# Demographic features and mortality risks in smallholder poultry farms of the Mekong river delta region: Poultry management and mortality risk

**DOI:** 10.1101/341800

**Authors:** Alexis Delabouglise, Benjamin Nguyen-Van-Yen, Nguyen Thi Le Thanh, Huynh Thi Ai Xuyen, Phuong Ngoc Tuyet, Ha Minh Lam, Maciej F Boni

## Abstract

This study describes the demographic structure and dynamics of small scale poultry farms of the Mekong river delta region, one of the world’s highest-risk regions for avian influenza outbreaks. Fifty farms were monitored over a 20-month period, with farm sizes, species, age, arrival/departure of poultry, and farm management practices recorded monthly. The history of poultry flocks in the sampled farms was recovered using a flock-matching algorithm. Median flock population sizes were 16 for chickens (IQR: 10 – 40), 32 for ducks (IQR: 18 – 101) and 11 for Muscovy ducks (IQR: 7 – 18); farm size distributions for the three species were heavily right-skewed. There was substantial flock overlap on almost all farms, with only one farm practicing an all-in-all-out management system. The rate of interspecific contacts was high, with two out of three farms housing at least two bird species. Among poultry species, demographic dynamics varied. Muscovy ducks were kept for long periods, in small numbers and outdoors, while chickens and ducks were farmed in larger numbers, indoors or in pens, with more rapid flock turnover. Most chicks were sold young to be fattened on other farms, and broiler and layer ducks had a short production period and higher degree of specialization. The rate of mortality due to disease did not differ much among species, with birds being less likely to die from disease at older ages, but frequency of disease symptoms differed by species. Time series of disease-associated mortality and population size were correlated for Muscovy ducks (Kendall’s coefficient τ = 0.49, p value < 0.01).

**Impacts:**

- The structure and dynamics of poultry populations kept in small scale farms in the Mekong river delta were accurately described.
- Poultry farms mix poultry of different species, ages, and production types, with multiple overlapping flocks present on a farm at all times. This promotes the persistence of pathogens.
- The three main farmed poultry species of poultry (chickens, ducks and Muscovy ducks) are managed differently and have different demographic characteristics.

## Introduction

Avian influenza (AI) infections caused by the H5N1 and H7N9 subtypes have been among the most concerning emerging diseases of the past two decades (Li et al., 2004) and still constitute a major threat to both human and animal health (Lai et al., 2016). Since 2003, H5N1 has been endemic in the domestic poultry populations of several countries of South and Southeast Asia, including China, Vietnam, Indonesia and Bangladesh ( FAO, 2011). Endemic areas are mainly characterized by high densities of domestic poultry and free-grazing duck farming on flooded areas (Gilbert & Pfeiffer, 2012; Gilbert et al., 2008). National-level interventions to control the disease were centered on the development of surveillance systems, preventive culling of poultry in outbreak areas, temporary closure of live bird markets and, in the case of China, Hong Kong and Vietnam, mass poultry vaccination (Ellis et al., 2006; FAO, 2011; Leung et al., 2012; Sims, 2007). However, the epidemiology of the disease in domestic birds remains poorly understood, and as a result little is known of the true effectiveness of these control policies. For this reason, in-depth information on the poultry population dynamics and inter-species contacts, in areas where the disease is endemic, are needed.

Vietnam is one of the countries where H5N1 AI has remained endemic since it was first reported in 2003 (Delabouglise et al., 2017; Pfeiffer et al., 2007). Most of Vietnam’s domestic poultry population is farmed on a small scale level (<100 birds/farm) by a large number of rural households (>7 million) who practice poultry farming as a secondary activity and with limited financial investment and infrastructure (General Statistics Office of Vietnam, 2017a; Henning et al., 2013; Hong Hanh et al., 2007). Despite the recent development of a large-scale commercial sector, Vietnamese consumers display a strong preference for local breeds of chickens, which are mostly produced by smallholder farms (Hong Hanh, et al., 2007; Ifft, 2010). The role played by small scale poultry production systems in perpetuating the endemicity of the disease has been debated (Desvaux et al., 2011; K. A. Henning et al., 2009; Hong Hanh, et al., 2007; Thanh et al., 2017). On the one hand, smallholder farms are believed to have very limited biosecurity practices and often host several poultry species (most commonly chickens and ducks), which increases their susceptibility to AI transmission. On the other hand, their small size and slow turnover (rate of birth/introduction and sale/slaughter of poultry) may limit their capacity to amplify and sustain virus circulation for a long time. In addition, these farms are less well connected to live birds trade networks compared to larger commercial farms, which limits their capacity to spread the virus over long distances (Tung & Costales, 2007). For this reason, assessing population structure, demographic dynamics, and biosecurity practices of smallholder poultry farms is crucial. While poultry trade networks were investigated in several studies in northern Vietnam (Fournié et al., 2016) and farming practices of large itinerant duck flocks were studied in the south (Minh et al., 2010), little attention has been given to the specific management of small scale farms.

The present study aimed to collect descriptive data on the on-farm demographic structure and dynamics of poultry kept in small scale farms of Ca Mau province, located in the Mekong river delta region of Vietnam. The Mekong river delta hosts about one fifth of Vietnam’s domestic poultry (General Statistics Office of Vietnam, 2017b) and is one of the two main at-risk areas for H5N1 AI, due to its high density of domestic waterfowl and human population and its large periodically flooded areas (Gilbert et al., 2008; J. Henning et al., 2009).

## Materials and Methods

### 1. Data collection

As described in a previous publication (Thanh, et al., 2017), 50 smallholder poultry farms were enrolled in a longitudinal observational study in Ca Mau province at the southern tip of Vietnam. Three additional farms were enrolled during the course of the study after three farms discontinued participation. The study was carried out with the support and collaboration of the Ca Mau sub-Department of Livestock Production and Animal Health (CM-LPAH). The collaboration between the investigators (authors) and CM-LPAH was approved by The Hospital for Tropical Diseases in Ho Chi Minh City, Vietnam. The CM-LPAH specifically approved this study; at the province-level in Vietnam, CM-LPAH is the equivalent of an Animal Care and Use Committee that approves studies involving biological sampling from animals.

Study duration was 20 months, from June 2015 to January 2017. A Vietnamese-language farm questionnaire was collected at the end of each month of the study. The collected information included: number of birds of each species present on the farm and their production type (broiler or layer/breeder); expected age of removal from the farm; number of birds introduced, removed and deceased in the last month with, in case of death, associated cause and/or clinical symptoms; vaccines administered to birds with date of vaccination; type of housing and biosecurity practices applied in managing the birds. To facilitate data collection, each farm’s poultry were classified into groups (hereafter referred to as “flocks”) with the same age, species, and production type, and data were collected at the flock level rather than individual level; the flock is the natural unit of management for a Vietnamese smallholder poultry farmer. Questionnaires are available in **Supplementary Information 1**.

### 2. Data processing

The questionnaire data were input and stored in a SQLite relational database. To convert the cross-sectional dataset into a longitudinal dataset, the flocks labeled in monthly questionnaires needed to be identified as being the same flock or not (e.g. flock #3 in March with 40 2-month old chickens and flock #7 in April that had 40 3-month old chickens would be identified as the same flock). To perform this “flock matching”, we used data on flock age, species, production type, inflows (buying and hatching), and outflows (selling, slaughtering, and other causes of death), ensuring that other flock characteristics were as similar as possible. These matches were performed on a per-farm basis, looking at two consecutive monthly questionnaires at a time.

With approximately 1000 questionnaires to process, we developed a flock-matching algorithm to process the data, and we checked consistency and correctness manually. From one month to the next, each flock must have at most one match, but flocks can also appear (buying and hatching) or disappear (selling, slaughtering). We want the state of a flock to correspond as much as possible to the expected state of its match in the previous month. This problem is a variant of an assignment problem for bipartite graphs, the goal being to minimize an objective function, for any farm at any month:

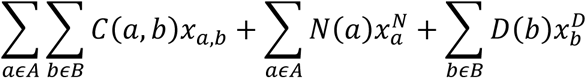

where *A* is the set of flocks at month *i, B* is the set of flocks on the same farm at month *i-1. x_a,b_* is 1 if the flock a is paired with the flock *b*, and 0 otherwise. *C*(*a*, *b*) is the cost of pairing *a* to *b*, (see Supplementary Information 2). *N(a)* is the cost of making *a* a new flock (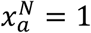), and *D(b)* the cost of making *b* a removed flock (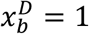). The objective function must be minimized under a set of constraints, to ensure that each flock has at most one match:

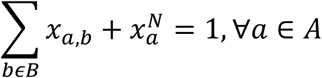

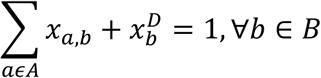

The algorithm is implemented in Python (v3.0), and is available online (Nguyen Van Yen, 2017). More details on the algorithm and the cost functions used are available in the **Supplementary Information 2**.

### 3. Data analysis

Rates of poultry removal and death were estimated. As the number of deaths and removals were collected on a monthly basis, the exact numbers of birds introduced, removed, and deceased on each day were not available. Thus, the day of introduction/removal/death was imputed assuming a uniform probability of this event occurring at any day during the month, and 10,000 imputed data sets (with exact days of birth/death events) were used to provide estimates and ranges of death rates and removal rates in the poultry population. Different probability distributions (exponential and mixtures of one, two, and three gamma distributions) were fit to the distributions of flock sizes, where the flock size is defined as its maximum size during its existence. Best fits were determined through maximum-likelihood estimation, and in the case of mixtures, using the expectation maximization algorithm of the mixtool R package (Benaglia, Chauveau, Hunter, & Young, 2009). Best fits were chosen according to Akaike Information Criterion (AIC). All data analyses were performed using R version 3.3 (R core team, 2014).

## Results

From June 2015 to January 2017, a total of 53 poultry farms were monitored, 26 in Tan Loc commune and 27 in Tan Phu commune. The questionnaire data during this period described 1087 discrete poultry flocks comprising 110,232 birds: 48,356 chickens, 33,570 quails, 25,450 ducks, 2443 Muscovy ducks, 195 geese, 183 pheasants, and 35 pigeons. Chickens, ducks and Muscovy ducks (MD) were the most common species on the farms, and these are known to be the three common types of poultry raised, sold, and eaten in southern Vietnam (Desvaux, Ton, Thang, & Hoa, 2008); quails were present on four study farms. Flock timelines are shown in **Supplementary Information 3** (plots per farm) and **Supplementary Information 4** (plots per commune). Only one of the monitored farms implemented all-in-all-out management throughout the study period. There was substantial flock overlap on all others farms.

### 1. Distribution of flock sizes and species

The distribution of the number of chickens and ducks per flock was highly right-skewed and over-dispersed; the mean flock sizes were 40 (chicken), 81 (ducks) and 14 (MD) while the median flock sizes were 16 for chickens (inter-quartile range (IQR): 10 – 40), 32 for ducks (IQR: 18 – 101) and 11 for Muscovy ducks (IQR: 7 – 18). A mixture of two or three gamma components gave the best fit for the distribution of flock sizes (three components for chickens and Muscovy ducks, two components for ducks) (**Supplementary Information 5**).

Only four farmers kept a single species of poultry during the study period. At any given time, poultry farms had a 32% probability of containing birds of only one species, a 33% probability of having birds of two different species, and a 35% probability of having birds of at least three different species. For a given species, the probability that it would be in contact with poultry of another species, on the same farm in the same month, is shown in **Table 1**. The frequency of contact between birds of different species was high, especially among the three most common species. In farms combining at least two of the three main species, the correlation between the numbers of birds of each pair of species was assessed using Kendall’s rank correlation coefficient. The number of ducks was positively correlated with the number of MDs (*τ* = 0.14, *p*-value <0.01) and chickens (*τ* = 0.07, *p*-value =0.042). The correlation between the number of chickens and MDs was not significant (*p*-value =0.62).

**Table 1.**
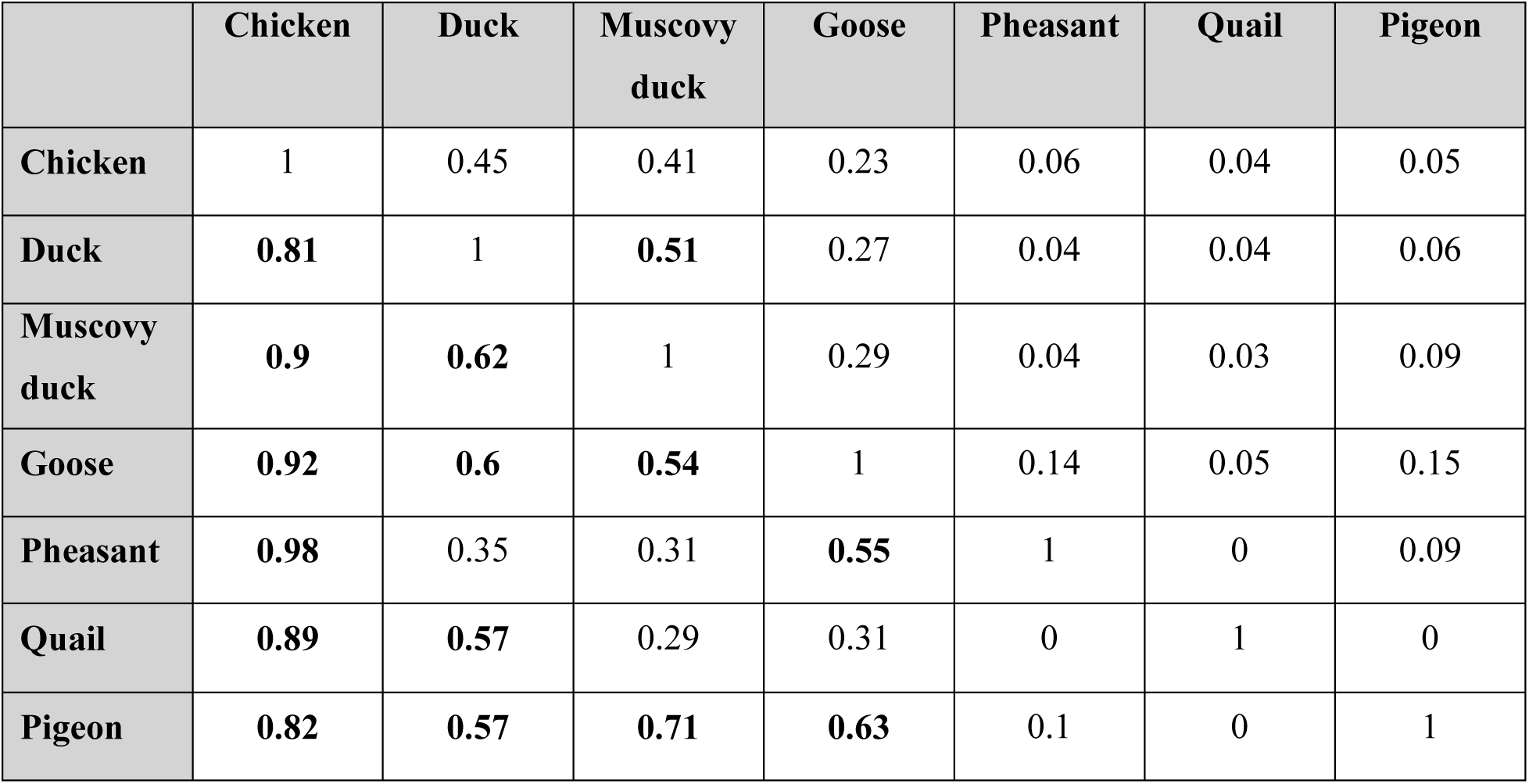
Interspecific interaction matrix on poultry farms. The probability that a bird of one species (row) will be in contact with birds of another species (column).

### 2. Life cycle of poultry

Distribution of age of departure from the farms and simulated distributions (from imputed data) of rates of removal and death due to disease according to poultry age in the three main poultry species are shown in **Figure 2**. Three stages of production were distinguished: young (chicks or ducklings, <1month old), broiler (>1 month old, mainly grown to be slaughtered for meat production), and layer/breeders (>1 month old, females kept for egg production and/or reproduction).

**Figure 1.**
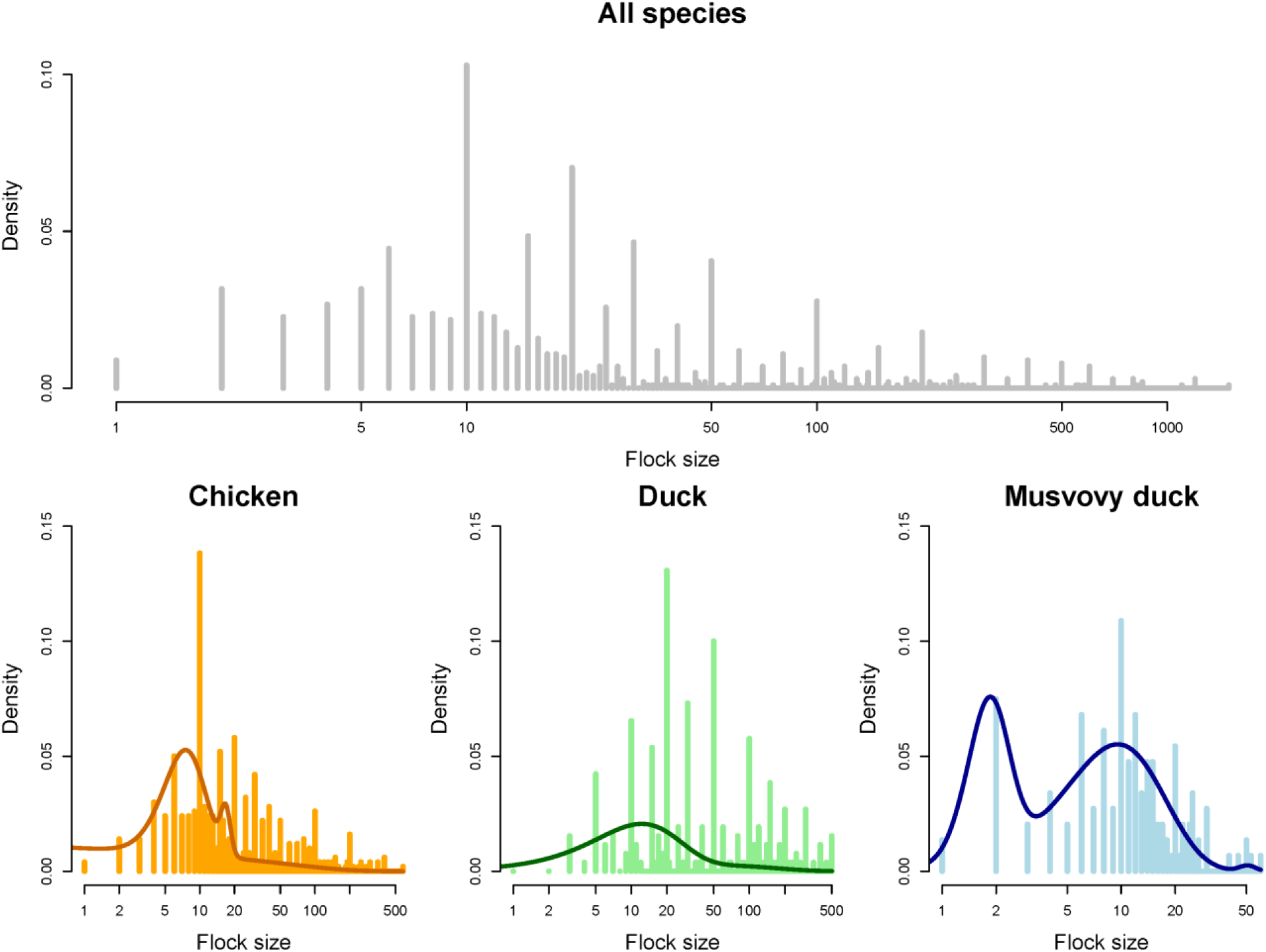
Distribution of the sizes of poultry flocks (Top, all species aggregated; bottom, plotted by species) in the studied population. Solid lines represent best-fit distributions (mixture of two gamma components (ducks) or three gamma components (chickens and Muscovy ducks)).

**Figure 2.**
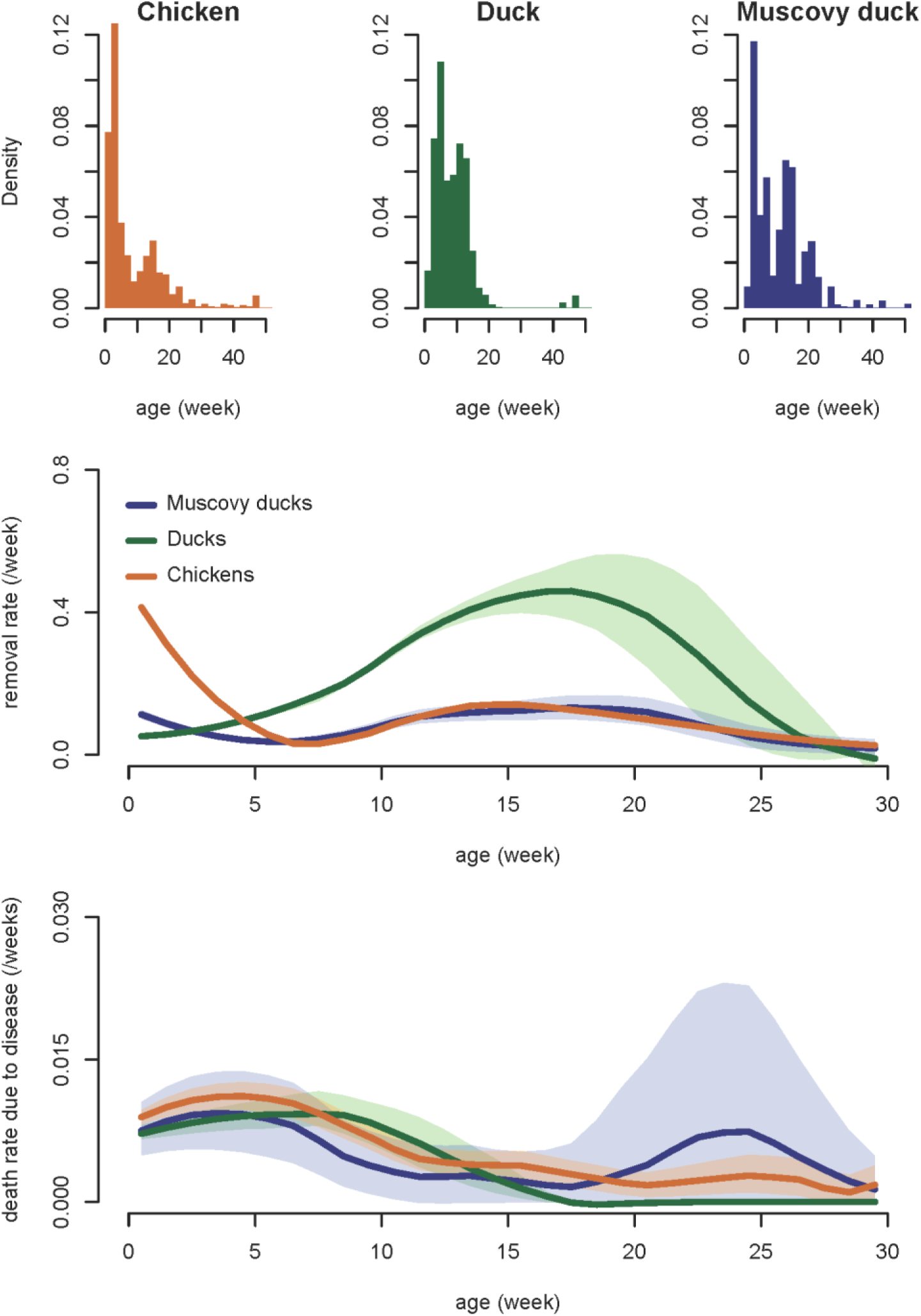
Age-related mortality and removal in the three main species of the studied poultry population. **Top:** distribution of ages at departure (death or removal). **Middle:** rate of removal as a function of age. **Bottom:** disease-related death rate as a function of age. Shaded regions in the middle and bottom graphs show the 95% range using imputed values; solid lines show the median. The time series of median, 2.5%, and 97.5% quantiles were smoothed using local regression (span factor: 0.5).

The age-specific removal rate was consistently higher for ducks than for chickens or MDs, except during the first 3 weeks, while the removal rate of MDs was the lowest (**Figure 2, middle**). 70% of young chickens, 45% of young ducks and 38% of young MDs were removed from the farm before reaching their first month. The high removal rate of young chickens, and, to a lesser extent, of MDs, was partly attributable to the presence of poultry farms with a high breeding activity (breeding and sale of young birds to be grown on other farms) while most sold young ducks were used to feed pythons. This explains the two clusters of age at departure in chickens, one corresponding to sale of newborn chicks, the other to the sale of mature broilers (**Figure 2, top**). For all three species, more than 90% of adult broilers came from the young stock of the farm. On average broiler chickens and MDs were kept for 12-14 weeks and broiler ducks for 7 weeks (**Figure 3**).

**Figure 3.**
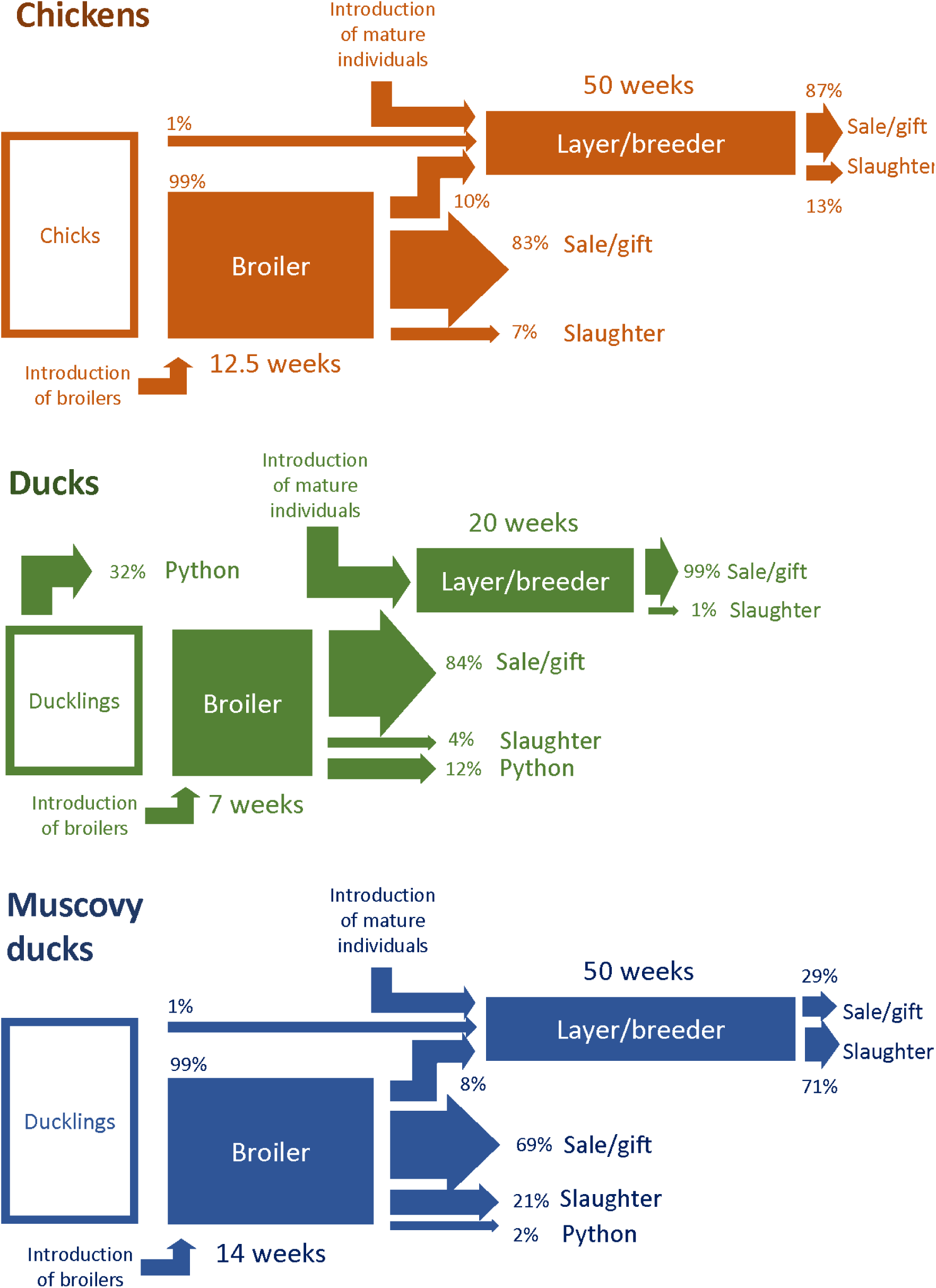
Flow diagram representation of the life history of poultry grown in the studied population.

Layer/breeder poultry were much older than broilers (average age above one year in the three main species). The ratio of layers/breeders to broilers was twice as high in chickens and MDs (0.41 in both species) than in ducks (0.20). In addition, the ratio of layer/breeder per introduction (addition of a new bird on a farm) was lower in ducks (0.19) than in chickens (0.37) and MDs (0.66). All layer/breeders ducks were introduced from another farm at a mature age while 63% of layer/breeder chickens and 68% of layer/breeder MDs were previously raised in the same farm, most of them being first part of a broiler flock and later kept for egg production or breeding instead of being sold or slaughtered (**Figure 3**).

The difference in the modality of supply of layer/breeder individuals was translated in the distribution of age at departure. In ducks, most individuals departed before 20 weeks of age (broilers), and all others after 40 weeks (layer/breeders), while in the two other species some birds departed in between these two age ranges (**Figure 2**). The relatively high removal rate of chicks and relatively short production period of adult ducks translated into a high rate of bird introduction (number of introductions of new individuals per month divided by the population size) which was twice as high in chickens and ducks (0.66 and 0.62 respectively) than in MDs (0.36).

### 3. Temporal population dynamics

Time series of population size and mortality rate attributable to disease are shown in **Figure 4**. Chicken population size slightly decreased during the study period and was not obviously seasonal. Duck population size peaked in October while the population of MDs increased at the end of each calendar year.

**Figure 4.**
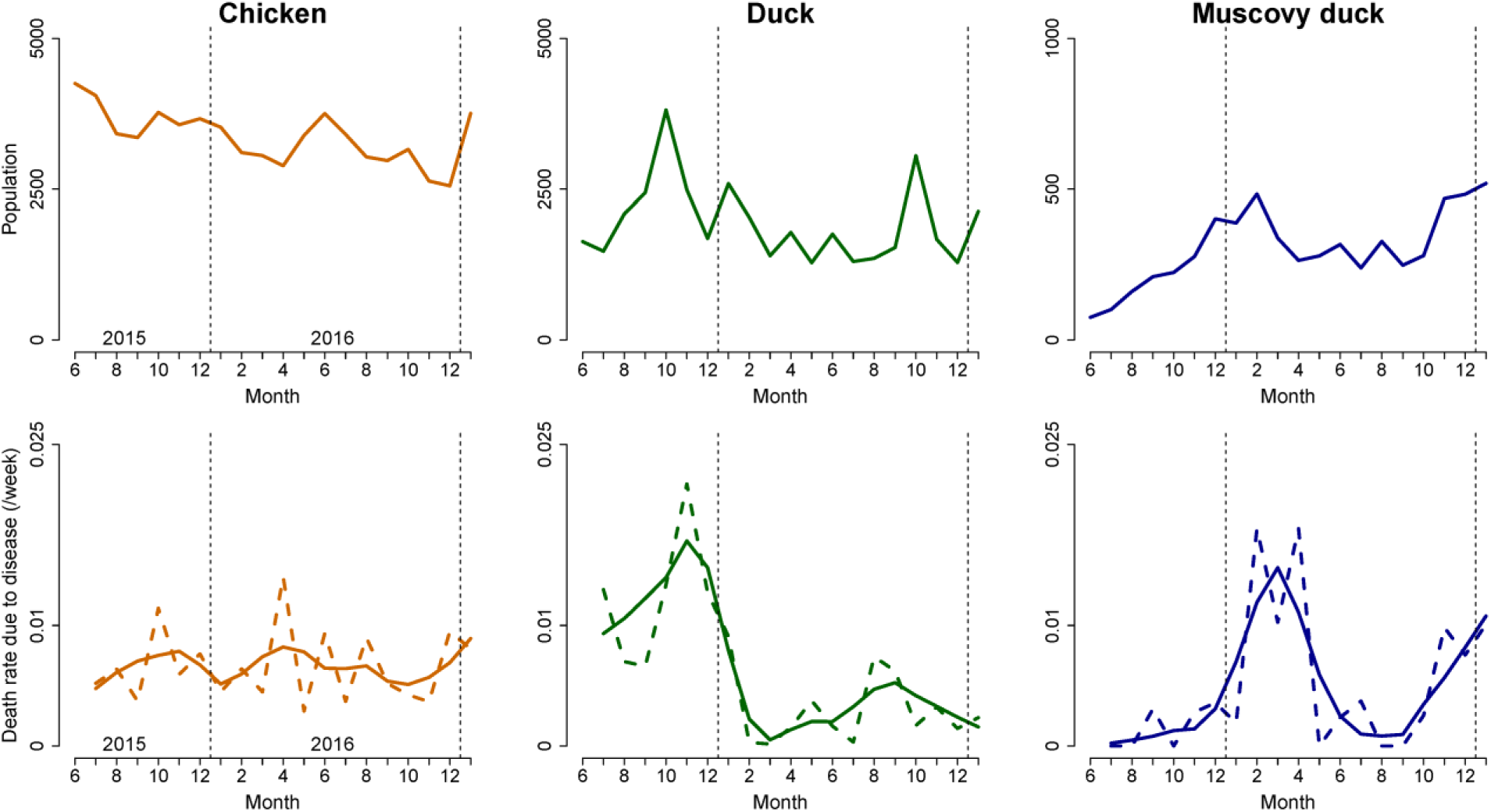
Evolution of population size and mortality rate attributable to disease over the study period. Bottom: Dashed lines are median estimates of death rates while solid lines are smoothed rates obtained through local regression (span factor: 0.5).

### 4. Poultry mortality

8.2% of birds died during the course of the study instead of being sold, given, slaughtered or fed to pythons. 44% of these deaths could be associated with a disease state (observation of clinical symptoms by farmers, diagnosis of a specific disease by farmers, or sudden death of several birds with no obvious cause). Other causes of death were accidents (disappearance of birds during grazing period, injuries from fighting, or attacks by rats or snakes) (1.2%), cold temperatures (0.4%), and unidentified reasons (3.0%). Note that these figures probably underestimate the true amount of loss due to disease since a few farmers explicitly mentioned they fed their sick poultry to pythons instead of letting them die. Poultry farms had a 4.4% risk of facing a mortality rate due to disease of more than 20% in a single month. Such events are indicated on the farm timelines in **Supplementary Information 3**.

Frequencies of observation of clinical symptoms when disease-related deaths occurred are shown in **Figure 5**. Poultry deaths were often associated with lethargy/weariness and digestive symptoms in the three species. Respiratory symptoms were specifically often reported in chickens while leg paralysis and nervous-system symptoms were quite specific to ducks. The mortality rate attributable to disease appears to be a decreasing function of age (**Figure 3**), approximately 1% per week in birds below 5 weeks of age and decreasing progressively afterwards. In MDs, however, an increase in mortality rate attributable to disease was observed around 20 weeks of age.

**Figure 5.**
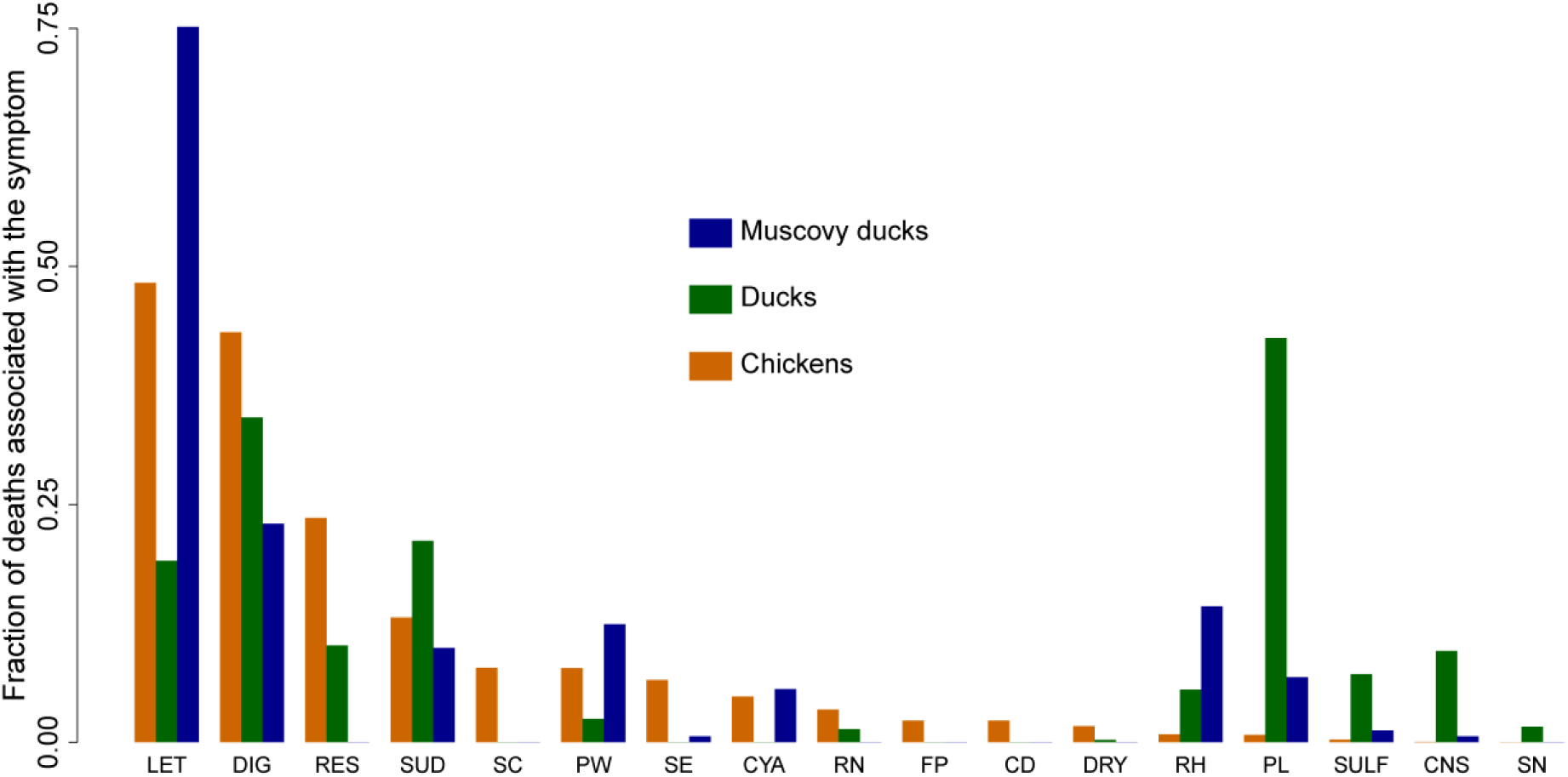
Frequency of association of poultry deaths attributable to diseases with specific symptoms or suspected causes in the studied poultry population. LET: lethargy, weariness; DIG: digestive symptom (diarrhea, flatulence or abnormal color of feces); RES: symptoms related to the lower respiratory tract (dyspnea or increase respiratory sounds); SUD: sudden death (birds died before any symptoms could be noticed); SC: swollen crop; PW: paralyzed wing; SE: anorexia; CYA: cyanosis; RN: symptoms related to the upper respiratory tract (runny nose); FP: fowl pox; CD: coccidiosis; DRY: dry legs; RH: retraction of the head; PL: paralyzed leg; SULF: intoxication with sulfate; CNS: nervous-system symptoms; SN: shrinking neck.

In chickens, the rate of mortality attributable to disease did not vary much across the study period, except for a peak in April 2016. In ducks, two peaks of mortality attributable to disease occurred during the study period. The major one (in November-December 2015), with 10% monthly mortality, seemingly followed a peak in duck population (**Figure 4**). In MDs two peaks in disease-related mortality (from around 1%/month to more than 5%/month) were observed in early and late 2016. Both increases seemed to follow a peak in the MDs’ population size. Time series of median estimates of disease-associated mortality rate and population size at the beginning of the month were significantly positively correlated for Muscovy ducks (Kendall’s coefficient τ = 0.49, *p*-value < 0.01). In chickens and ducks the correlation was not significant (*p*-values of 0.63 and 0.53, respectively).

### 5. Housing and grazing of poultry and infectious disease prevention

Approximately half of chickens were housed indoors while the overwhelming majority of ducks and MDs were farmed outdoors, either in pens (ducks) or free-range (most MDs) (**Figure 6**). 69% of ducks grazed in water bodies during the day. The average grazing distance was 108m away from the farm and 1.5 km was the maximum.

**Figure 6.**
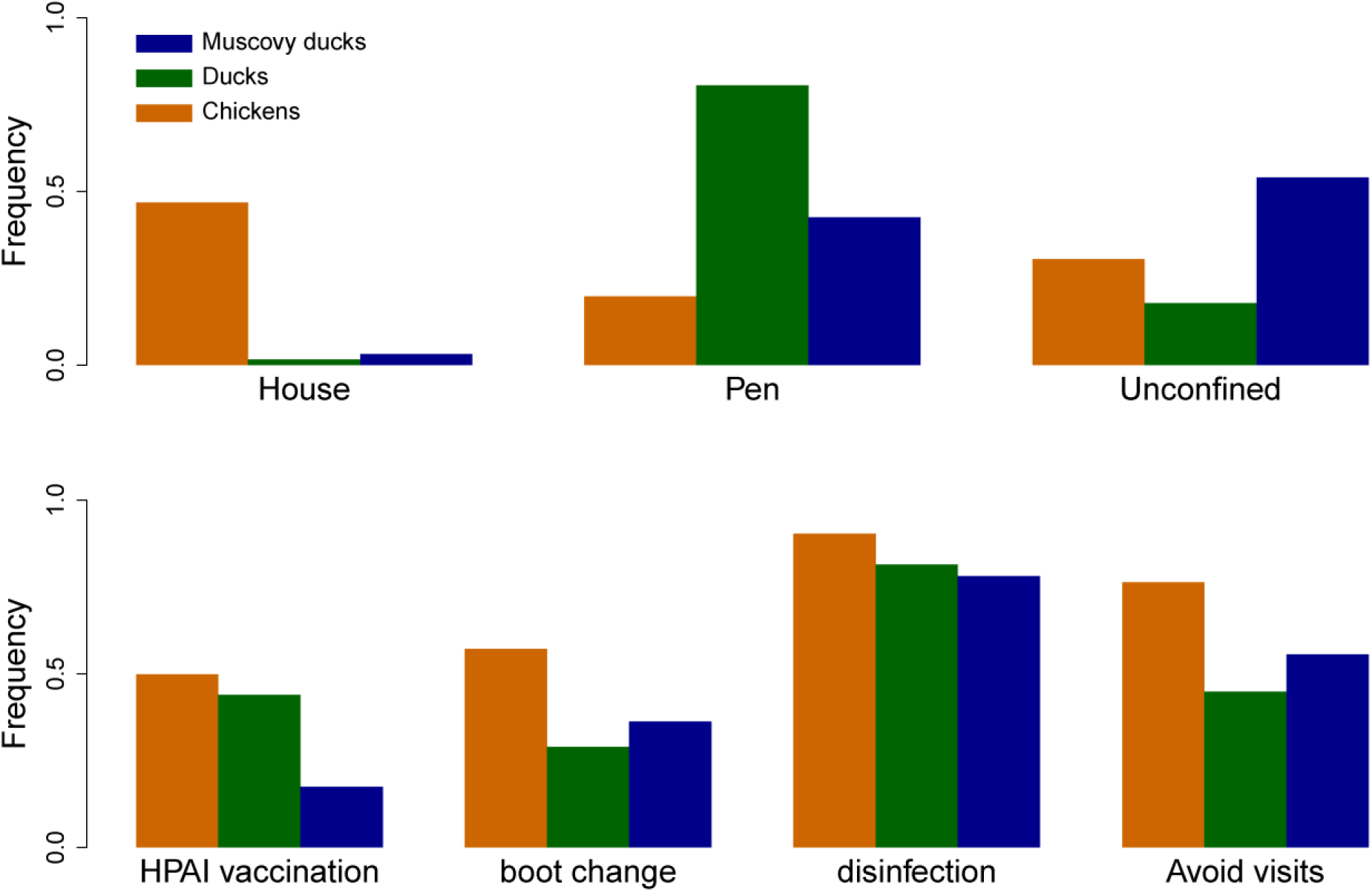
Type of housing and frequency of application of different prevention practices against infectious diseases in the studied poultry population.

The vaccination coverage against AI was 50% in chickens, 44% in ducks and 17% in MDs. The vaccination coverage against Newcastle Disease was 26% in chickens while vaccination coverage against duck plague was 1.5% in ducks. The other diseases against which vaccination was practiced were Fowl cholera and Gumboro disease, mainly in chickens.

## Discussion

Assessments of the demographic structure and dynamics of small scale poultry farms are made difficult by the lack of systematic accounting of birds’ entries and exits on farms, an absence of all-in-all-out management, simultaneous presence of birds of different species and production stages on the same farm, and combination of self-renewal and outsourcing of young birds from other farms and/or hatcheries. Longitudinal surveys, such as the one conducted in this study, are therefore necessary. One difficulty in such longitudinal surveys is the tracing of each identified birds from one month to the next. This difficulty was overcome through a flock-matching algorithm. The study data combined with the flock-matching algorithm produced an accurate description of smallholder farms’ demographic structure and dynamics in the Mekong river delta region.

All-in-all-out flock management was nearly absent and multispecies farming was widespread. Farmers usually kept at least two different species of birds, and chickens had an almost 50% chance to be farmed together with ducks. The on-farm number of ducks was positively correlated with the number of Muscovy ducks and chickens. In other studies, multispecies poultry farms have been shown to be at higher risk of introduction of AI (Desvaux et al., 2011; Henning et al., 2011). However, in the context of small scale farming, with highly fluctuating input and output market prices and absent or limited risk-mitigation mechanisms (contracts with suppliers or buyers, insurance schemes and/or vertical integration) (ACI, 2006; Burgos et al., 2008; Delabouglise et al., 2016; Tung & Costales, 2007), diversification of livestock production constitutes a way of limiting income variation over time. In addition, duck farming is deeply integrated in the rice production systems of the Mekong region since scavenging ducks feed on grains left in freshly harvested rice fields and, in the same time, reduce the density of parasite snails (Minh, et al., 2010).

Farmers lost at least 3.6% of their production to disease. In Muscovy ducks, major peaks of disease-related mortality closely followed peaks in population size. The same visual association could be seen for ducks but its statistical significance could not be demonstrated. The chicken population size and disease-attributed mortality had less temporal variation and did not appear to correlate temporally with each other. These observations are consistent with the hypothesis that the transmission of infectious diseases in smallholder domestic poultry farms has a density-dependent component and, therefore, that any increase in the domestic bird population puts farms at higher risk of disease-induced losses (Boni et al., 2013; Otte et al., 2008). Variation in poultry population can be driven by changes in demand for poultry products, as occurs during the Lunar New Year festival (Delabouglise et al., 2017), or by changes in environmental conditions. For instance, peaks in duck population in October are probably related to rice production cycles which are quite specific to each area of the Mekong river delta region (Minh et al., 2010). The lack of observed seasonality in the chicken population is surprising, considering the observed increase in demand for chicken meat at Lunar New Year. Nonetheless this increase in demand is reportedly less pronounced in the southernmost provinces of Vietnam (Delabouglise et al., 2017).

This study highlights some species-specific farm characteristics that may influence the epidemiology of infectious diseases in poultry. Muscovy ducks are kept outdoors, mostly unconfined, and vaccination uptake is very low. MDs may be more at risk of contact with pathogens released in the environment or with infectious birds sharing the same scavenging area (K.A. Henning et al., 2009), and the epidemiology of disease transmission in MDs is likely to be strongly influenced by the individual farm that they are housed on. Chickens and ducks, on the other hand, show a much higher replacement rate (the rate of removal and introduction of birds). Thus, chicken and duck farms may be more at risk of introduction of pathogens through inter-farm movements, and the epidemiology of avian influenza transmission in chickens and ducks may be more strongly influenced by the trading network than by individual farm characteristics.

The results suggest different modalities of renewal of the poultry stock in the three species. In chickens, a substantial number of smallholder farms bred and supplied other smallholder farms with chicks, which explains the high removal rate of chickens in their two first weeks of life (>40%/week). Since the origin of the young birds (bred and hatched in the farm or supplied from other farms or hatcheries) was not informed in the questionnaires, it is unclear to which extent the studied poultry farms practice self-renewal or outsource their young stock from outside. However, it is evident that the sale of ducklings to supply other smallholder farms is more limited. The low ratio of layer/breeder per poultry introduction in ducks tends to suggest that a significant fraction of ducklings are purchased from other types of suppliers (large breeding farms and hatcheries). Conversely, the high ratio of layer/breeders per poultry introduction in MDs tends to suggest MD farms maintain their population essentially through self-renewal (birds are bred, hatched and grown in the same farm).

The three main poultry species have markedly different turnovers. In ducks the production period of both broiler and layers is short. In addition, no layer ducks are bred and raised on the same farm, broilers are never converted in layer/breeders, and the ratio breeder/layer over bird introduction is particularly low, suggesting a high rate of import of new individuals from outside. These observations are important, considering the major role played by ducks, and particularly broiler ducks in the epidemiology of AI in the Mekong river delta (Henning et al., 2011; Nguyen et al., 2014). They can be explained by a higher degree production efficiency and specialization of the used breeds, shortening their growth period, and increasing the outsourcing of broilers and layers from breeding farms. According to previous studies, duck farms seem to use a high proportion of exotic breeds while small chicken farms tend to use preferentially indigenous breeds, which are more appealing to Vietnamese consumers (Duc & Long, 2008; Henning et al., 2013; Ifft, 2010). Finally about 40% of ducks were fed to pythons (most at <1-month old). Python meat consumption is common in the Mekong river delta and the waste protein produced by the poultry industry is mentioned as an important source of feed (Aust, 2015).

Control of avian influenza in Asia will likely continue employing poultry vaccination, responsive depopulation, and basic farm management/biosecurity implementation tools as its main methods. A better understanding of poultry turnover rates, species overlap, flock overlap, species-specific infections risks, and the influence of environmental transmission and trader/market based transmission will be critical for designing control and response policies that can be tailored to regions with known poultry species compositions and known farm management practices.

## Supplementary Information

**Supplementary information 1.** Questionnaires in English and Vietnamese version

**Supplementary information 2.** Description of the flock-matching algorithm

**Supplementary information 3.** Timeline of poultry flocks in each study farm

**Supplementary information 4.** Timeline of poultry flocks in each study commune

**Supplementary information 5.** Characteristics of fitted probability distributions of flock sizes of the three main poultry species

## Acknowledgements

Many thanks to the field staff in Ca Mau, especially Mr Huynh Thanh Tuan and Mrs Duong Bich Ngan. The study was funded by the Defense Threats Reduction Agency (US) and by Wellcome Trust grant (098511/Z/12/Z). AD and MFB are funded by Pennsylvania State University. The funders had no role in study design, data collection and analysis, decision to publish, or preparation of the manuscript.

